# Characterization of Pik1 function in fission yeast reveals its conserved role in lipid synthesis and not cytokinesis

**DOI:** 10.1101/2023.07.24.550375

**Authors:** Alaina H. Willet, Lesley A. Turner, Joshua S. Park, Liping Ren, Chloe E. Snider, Kathleen L. Gould

## Abstract

Phosphatidylinositol (PI)-4-phosphate (PI4P) is a lipid found at the plasma membrane (PM) and Golgi in cells from yeast to humans. PI4P is generated from PI by PI4-kinases and can be converted to PI-4,5-bisphosphate [PI(4,5)P_2_]. *Schizosaccharomyces pombe* have 2 essential PI4-kinases: Stt4 and Pik1. Stt4 localizes to the PM and its loss from the PM results in a decrease of PM PI4P and PI(4,5)P_2_. As a result, cells divide non-medially due to disrupted cytokinetic ring-PM anchoring. However, the localization and function of *S. pombe* Pik1 has not been thoroughly examined. Here, we found that Pik1 localizes exclusively to the trans-Golgi and is required for Golgi PI4P production. We determined that Ncs1 regulates Pik1, but unlike in other organisms, it is not required for Pik1 Golgi localization. When Pik1 function was disrupted, PM PI4P but not PI(4,5)P_2_ levels were reduced, a major difference with Stt4. We conclude that Stt4 is the chief enzyme responsible for producing the PI4P that generates PI(4,5)P_2_. Also, that cells with disrupted Pik1 do not divide asymmetrically highlights the specific importance of PM PI(4,5)P_2_ for cytokinetic ring-PM anchoring.

**Summary statement:** Fission yeast Pik1 localizes exclusively to the trans-Golgi independently of Ncs1, where it contributes to PI4P but not PI(4,5)P_2_ synthesis. Pik1 does not affect cytokinesis.

## Introduction

Phosphoinositides (PIPs) are abundant lipid species important for a variety of cellular processes including cell division and membrane trafficking (Echard, 2012; Schuh and Audhya, 2012). Phosphatidylinositol (PI)-4-phosphate (PI4P), which is made by PI-4-kinases phosphorylating the head group of PI, is a precursor for PI-4,5-bisphosphate [PI(4,5)P_2_] and both lipid species are important for cell division in diverse organisms (Emoto *et al*., 2005; Field *et al*., 2005; Snider *et al*., 2017, 2018).

The fission yeast, *Schizosaccharomyces pombe*, has three PI-4-kinases: Stt4, Lsb6, and Pik1 and two PI4 phosphatases Sac11 and Sac12 (Fig. 1A). Stt4 is an essential type IIIɑ enzyme that localizes to the PM via two scaffolds, Efr3 and Ypp1 (Baird *et al*., 2008; Snider *et al*., 2017). Efr3 is critical for Stt4 PM localization and *efr3Δ* cells have reduced PM PI4P and PI(4,5)P_2_ (Snider *et al*., 2017). For *S. pombe* cells to divide, they build an actin- and myosin-based cytokinetic ring (CR) at the cell middle that guides the deposition of a medial septum (Marks *et al*., 1986; Kitayama *et al*., 1997). Normally the CR remains anchored at its original medial position so that upon CR constriction and septation, two daughter cells of equal size are produced (Snider *et al*., 2017). However, *efr3Δ* cells divide asymmetrically because the CR is not properly anchored and it slides toward one cell end prior to constriction and septation (Snider *et al*., 2017). It was hypothesized that PI(4,5)P_2_ is particularly important for CR anchoring however a role for PI4P in this process could not be ruled out (Snider *et al*., 2017, 2018).

**Figure 1.**
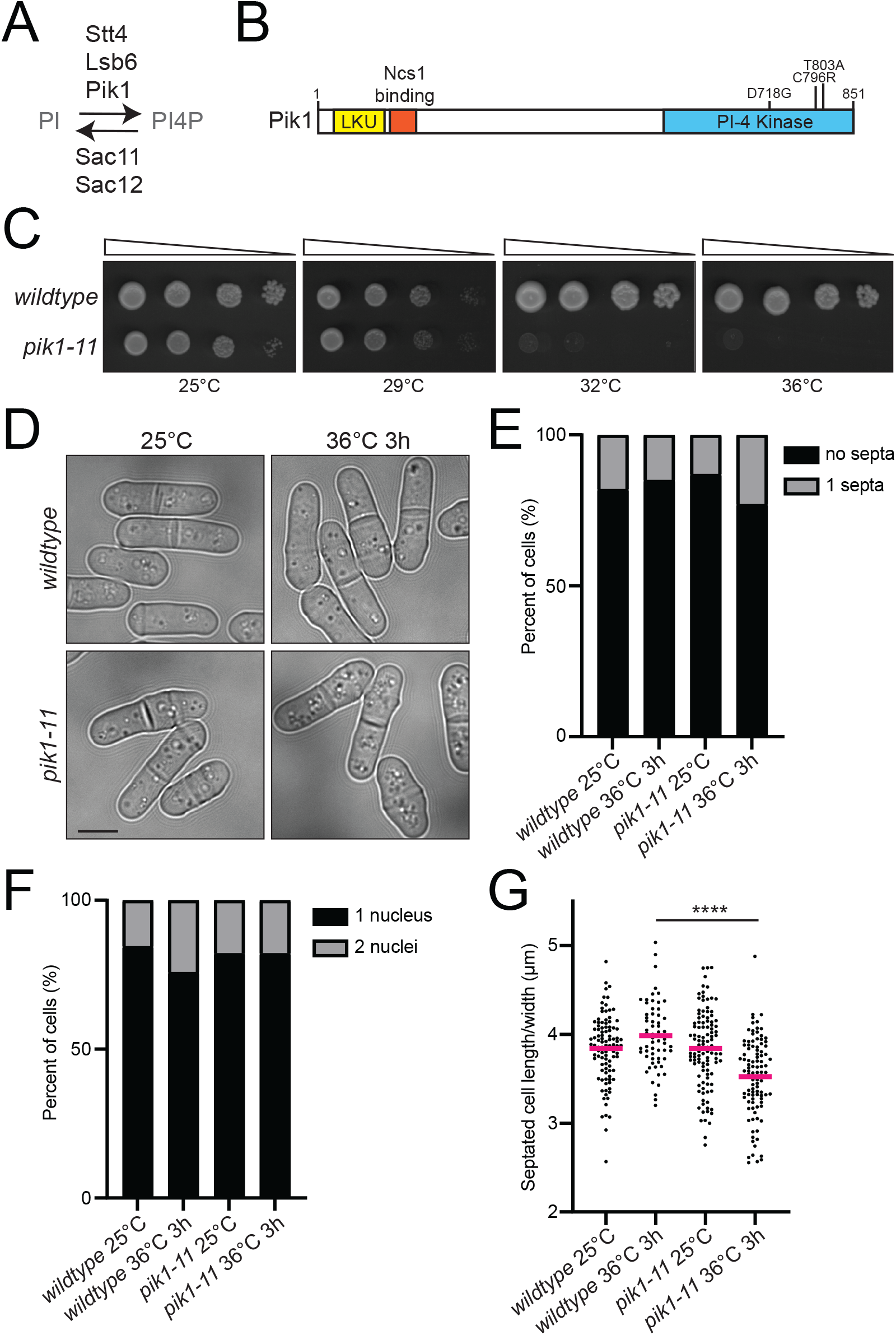
Isolation and characterization of *pik1-11*. A) A schematic of the *S. pombe* proteins known to phosphorylate PI to convert it to PI4P or to dephosphorylate PI4P. B) A schematic of Pik1, drawn to scale, with the domains and mutations encoded by the *pik1-11* allele labelled. LKU (Lipid Kinase Unique); PI-4 Kinase (phosphatidylinositol-4-kinase). C) 10-fold serial dilutions of the indicated strains grown at the indicated temperatures for 3-4 days on YE agar. D) Differential Interference Contrast (DIC) live-cell images of wildtype and *pik1-11* cells grown at 25°C and shifted or not to 36°C for 3 hours. Scale bar, 5 μm. E) Quantification of the septation index of the indicated strains at the indicated temperatures. n > 400 for each. F) Cells were grown up as in D and fixed and stained with DAPI and Methyl Blue (MB). Nuclei/cell were quantified. n ≥ 270 for each. G) Cells from A were measured for the ratio of septated cell length divided by the cell width. n ≥ 62 for each. **** *p* ≤ .0001. One-way ANOVA.

Lsb6 is a type II PI-4-kinase that localizes to the vacuolar membrane (Matsuyama *et al*., 2006). Cells lacking *lsb6* do not have any reported cellular defects (Kim *et al*., 2010), however they do have a reduction in PM PI4P and a negative genetic interaction with *efr3Δ* cells (Snider *et al*., 2018).

Lastly, Pik1 is an essential type IIIβ PI-4-kinase that has been extensively studied in *Saccharomyces cerevisiae*. In *S. cerevisiae*, Pik1 localizes to the Golgi, where it generates PI4P (Flanagan *et al*., 1993; Walch-Solimena and Novick, 1999) and is essential for proper Golgi to PM protein trafficking (Hama *et al*., 1999; Walch-Solimena and Novick, 1999). Pik1 complexes with the calcium binding protein Frq1 that is important for Pik1 kinase activity and Golgi localization (Hendricks *et al*., 1999; Ames *et al*., 2000; Strahl *et al*., 2005; Lim *et al*., 2011). Less is known about *S. pombe* Pik1. It is reported to localize to the Golgi, but also the cell division site, a localization not reported for Pik1 orthologs in other organisms (Park et al., 2009). *S. pombe* Pik1 was also reported to bind the Frq1 ortholog, Ncs1, which is thought to regulate its activity (Hamasaki-Katagiri *et al*., 2004). Interestingly, *S. pombe* Pik1 was also reported to be involved in cytokinesis and to bind the myosin light chains Cam2 and Cdc4 via a C-terminal pseudo isoleucine-glutamine (IQ) motif (Desautels *et al*., 2001; Sammons *et al*., 2011). These interactions are not reported for Pik1 orthologs in other species and the function of these potential interactions has been unclear.

Here, we aimed to clarify Pik1 function in *S. pombe* by constructing and characterizing a *pik1* temperature sensitive allele that we named *pik1-11* and constructing an endogenous fluorescently tagged *pik1* allele. Our results indicate that *S. pombe* Pik1 functions as the sole Golgi-localized PI4-kinase. Further, it does not contribute substantially to the PI4P precursors that are converted to PI(4,5)P_2_. Importantly, we found no evidence that *S. pombe* Pik1 co-localizes with Cdc4 or Cam2 or contributes to cytokinesis, but Ncs1 is indeed an important regulator of Pik1.

## Results and discussion

To investigate the function of Pik1, an essential PI4-kinase, we constructed a temperature sensitive allele that we named *pik1-11* (Fig.1B-C). The protein encoded by *pik1-11* contains three point mutations within the PI-4-kinase domain (Fig. 1B). *pik1-11* cells grow similarly to wildtype at 25°C and 29°C but *pik1-11* cells do not grow at 32°C or 36°C (Fig. 1C). A previously described *pik1* allele, *pik1-td*, displayed defects in cell division (Park *et al*., 2009). In contrast, we did not observe any difference in the septation index of *pik1-11* cells at permissive or restrictive temperatures compared to wildtype cells (Fig. 1D-E). We also did not observe an accumulation of multi-nucleated cells (Fig. 1F). Thus, we did not observe any cellular defect indicating a role for Pik1 in cell division. However, the morphology of *pik1-11* cells changed at the non-permissive temperature; they became shorter and wider (Fig. 1G).

Because Pik1 converts PI to PI4P, we investigated cellular PI4P levels with the established GFP-tagged lipid biosensor GFP-P4C_SidC_ (Luo *et al*., 2015; Snider *et al*., 2017). In wildtype cells at 25°C and after 3 hours at 36°C, GFP-P4C_SidC_ localized to the PM and internal puncta that correspond to the Golgi (Fig. 2A). GFP-P4C_SidC_ in *pik1-11* cells did not show any detectable Golgi localization and the PM intensity was reduced compared to wildtype at both permissive and restrictive temperatures (Fig. 2A-B). Because PI4P is a precursor of PI(4,5)P_2_, we next analyzed cellular PI(4,5)P_2_ localization with the GFP-2xPH_Plc_ lipid biosensor (Stefan *et al*., 2002; Snider *et al*., 2017). At 25°C, GFP-2xPH_Plc_ showed the expected PM localization in wildtype cells and in *pik1-11* cells there was no difference in cortical fluorescence intensity compared to wildtype (Fig. 2C-D). At 36°C, there was an ∼65% increase in cortical GFP-2xPH_Plc_ localization in *pik1-11* cells compared to wildtype (Fig. 2C-D). We conclude that *pik1-11* cells lack a Golgi PI4P pool, and have reduced PM PI4P that does not result in a corresponding decrease in PM PI(4,5)P_2_.

**Figure 2.**
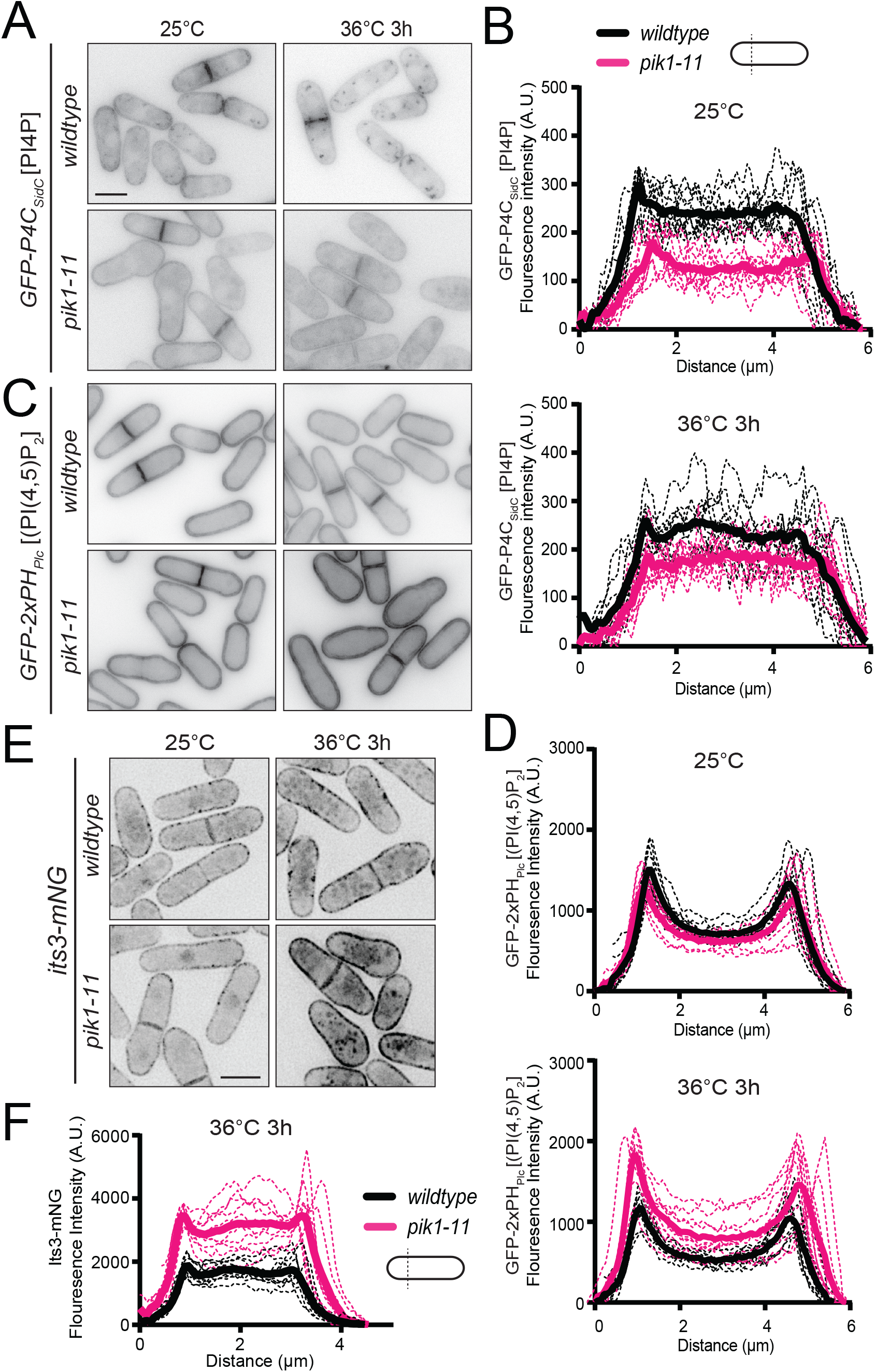
Analysis of lipid levels and septa position in *pik1-11*. Live-cell images of GFP-P4C_SidC_ (A) and GFP-2xPH_Plc_ (C) in wildtype and *pik1-11* cells. Cells were grown at 25°C and shifted or not to 36°C for 3 hours. B, D, F) Line scans of fluorescence intensity drawn across the short axis of 10 cells for each indicated strain at the indicated temperature. Solid lines represent the mean and dotted lines are the individual line scan traces. E) Live-cell images of Its3-mNG in wildtype and *pik1-11* cells. Cells were grown at 25°C and shifted to 36°C for 3 hours and imaged at both time points. Scale bars, 5 μm.

That *pik1-11* cells had reduced PM PI4P but not reduced PI(4,5)P_2_ was an intriguing observation. Other existing gene deletions and temperature sensitive alleles involved in PI4P synthesis (*efr3Δ*, *its3-1*) have reduced PM PI4P and also reduced PM PI(4,5)P_2_ (Snider *et al*., 2017, 2018) thus making dissecting the contributions of each lipid challenging. It was previously hypothesized that the CR anchoring defects of *efr3Δ* cells was most likely due to reduced PI(4,5)P_2_ rather than a change in PI4P because many CR proteins specifically bind PI(4,5)P_2_ (Cauvin and Echard, 2015; McDonald *et al*., 2015; Sun *et al*., 2015). The *pik1-11* allele allowed us to test this hypothesis more directly. We measured the septa position of wildtype and *pik1-11* cells at the permissive and restrictive temperatures and did not observe a difference in *pik1-11* compared to wildtype in any condition (Fig. S1A). This result is consistent with the hypothesis that altering PI4P levels alone do not impact CR anchoring to the PM. Similarly, *lsb6Δ* cells have reduced PM PI4P levels and do not have off-centered septa (Snider *et al*., 2018). However it was not reported if *lsb6Δ* cells have any changes in PM PI(4,5)P_2_ levels. To test this, we compared wildtype and *lsb6Δ* cells expressing GFP-2xPH_Plc_ and found no difference in the cortical fluorescence intensity of the PI(4,5)P_2_ sensor in these strains (Fig. S1B-C). Overall, these results suggest that Stt4, but not Pik1 or Lsb6, generates the PI4P at the PM that is subsequently converted to PI(4,5)P_2._ This conclusion is in accord with the exclusive localization of the PI4-5-kinase, Its3, to the PM (Zhang *et al*., 2000).

We next wanted to better understand why *pik1-11* cells have increased PM PI(4,5)P_2_ at the restrictive temperature. Given that disruption of *pik1* causes protein trafficking defects in *S. cerevisiae* (Hama *et al*., 1999; Walch-Solimena and Novick, 1999), we reasoned that *S. pombe* proteins involved in PI(4,5)P_2_ synthesis may be mis-localized. Specifically, an increase in PM Its3 or Stt4 and/or a decrease in PM PI4-5-phosphatases could account for the observed increased in PM PI(4,5)P_2_. We found no change in the localization of GFP-Stt4, or the PI4-5-phosphatases Inp53-mNG and Syj1-mNG (Snider *et al*., 2018) in *pik1-11* cells compared to wildtype at 25°C or 36°C (Fig. S1D). In contrast, PM Its3-mNG was mildly reduced at 25°C and increased 1.8-fold at 36°C in *pik1-11* compared to wildtype cells (Fig. 2E-F and S1E). We conclude that the increased PM PI(4,5)P_2_ and decreased PM PI4P observed in *pik1-11* cells at the restrictive temperature can be explained by additional Its3 activity at the PM.

We also tested if there were genetic interactions between *pik1-11* and other components of the PIP pathway. We did not observe a genetic interaction with *GFP-stt4*, a hypomorphic *stt4* allele (Snider *et al*., 2017) or *lsb6Δ* (Fig. S2). We did find a negative genetic interaction with *efr3Δ* and *its3-1*, but not with a deletion of *sac12*, a gene encoding a PI-4-phosphatase (Harris *et al*., 2022) (Fig. S2). Overall, we conclude that the PI4-kinases in *S. pombe* function independently, as in *S. cerevisiae* (Strahl *et al*., 2005), and disruption of other genes important for PI(4,5)P_2_ generation leads to further growth defects of *pik1-11* cells.

To better understand Pik1 function, we aimed to construct an endogenous fluorescently-tagged allele. We were not able to recover viable endogenous N- or C-terminally tagged *pik1* alleles. Therefore, we designed a construct that inserted mNG within a flexible loop of Pik1 based on the AlphaFold2 predicted structure (Jumper et al., 2021; Varadi et al., 2022) (Fig. 3A). Indeed, we were able to generate a *pik1-D450-mNG* strain, in which mNG was introduced after residue D450 in the endogenous *pik1* locus (Fig. 3A). This insertion is predicted not to disrupt the overall Pik1 fold (Jumper *et al*., 2021; Mirdita *et al*., 2022; Varadi *et al*., 2022) (Fig. 3A).

**Figure 3.**
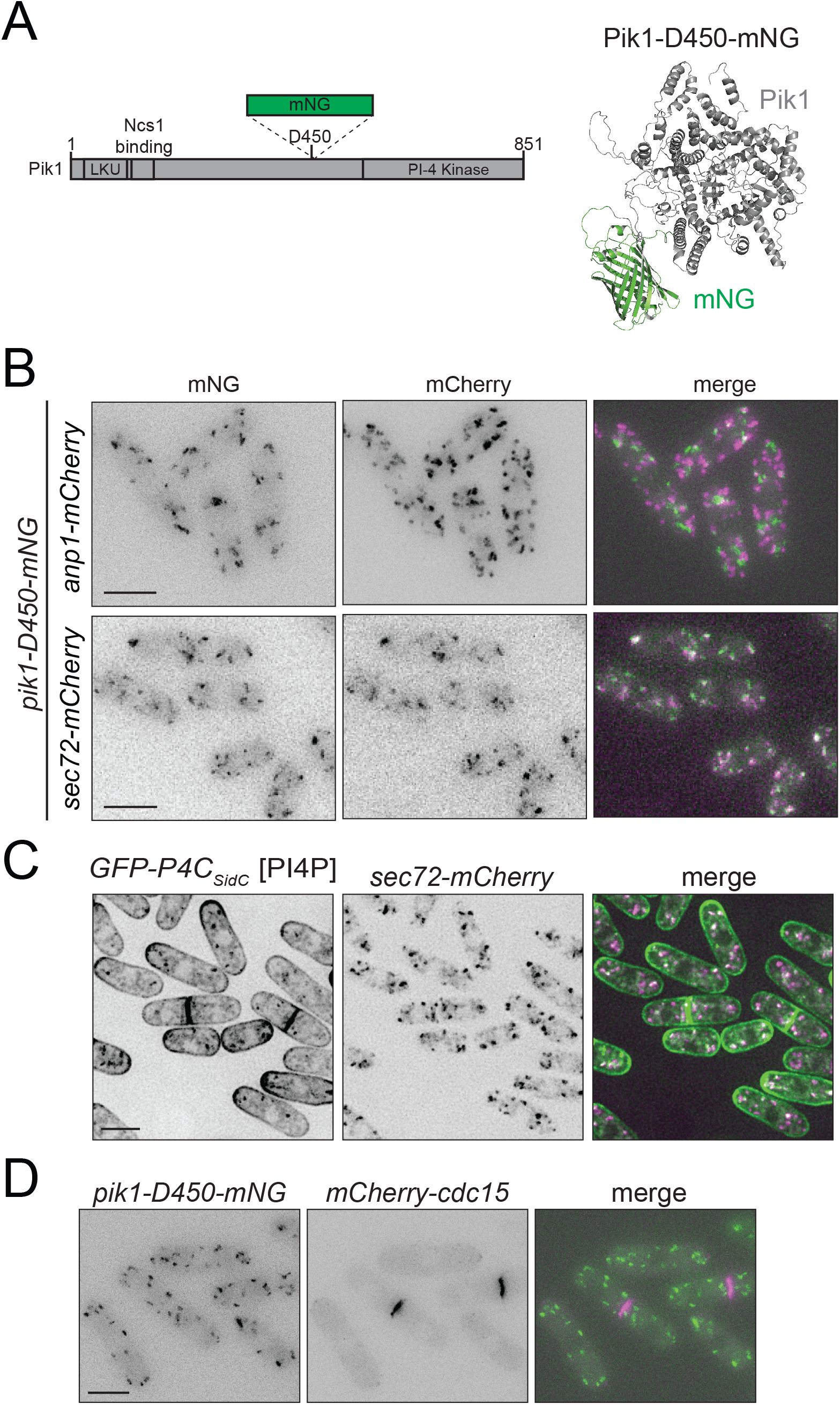
Pik1 localizes exclusively to the trans-Golgi. A) Left, a schematic of Pik1 drawn to scale with domains, mNG and the mNG insertion site labelled. Right, AlphaFold2 predicted structure of Pik1 with mNG inserted after residue D450. Pik1 is in gray and mNG is in green. B) Live-cell images of cells expressing Pik1-D450-mNG with either Anp1-mCherry or Sec72-mCherry. C) Live-cell imaging of cells expressing GFP-P4C_SidC_ with Sec72-mCherry. D) Live-cell imaging of cells expressing Pik1-D450-mNG with mCherry-Cdc15. Scale bars, 5 μm.

To investigate Pik1 localization, we co-imaged Pik1-D450-mNG with either a cis-(Anp1-mCherry) or trans-Golgi (Sec72-mCherry) marker (Vjestica *et al*., 2008) and found co-localization only with the trans-Golgi marker (Fig. 3B), consistent with *S. cerevisiae* Pik1 localization (Walch-Solimena and Novick, 1999; Sciorra *et al*., 2005; Strahl *et al*., 2005). We also found that the Golgi PI4P pool detected by GFP-P4C_SidC_ co-localized with Sec72-mCherry trans-Golgi marker, consistent with the Pik1 localization (Fig. 3C). We did not detect Pik1-D450-mNG at the cell division site when co-imaged with the CR marker, mCherry-Cdc15 (Fig. 3D). We conclude that Pik1 localizes exclusively to the trans-Golgi and that the previously reported localization to the division site might be explained as an artifact of over-expression (Park *et al*., 2009).

Additional evidence that Pik1 could be involved in cytokinesis is its reported interaction with the myosin light chains Cam2 and Cdc4 (Desautels *et al*., 2001; Sammons *et al*., 2011). When Pik1-D450-mNG was co-imaged with Cam2-mCherry, no co-localization was detected, consistent with Cam2 localizing exclusively to endocytic actin patches in wildtype cells (Fig. S3A) (Sammons *et al*., 2011). When released from its binding partner, myosin-1, with the *myo1ΔIQ* mutant, Cam2 localizes to non-actin patch puncta (Sammons *et al*., 2011) that we reasoned might be the Golgi. Thus, we imaged Pik1-D450-mNG and Cam2-mCherry in *myo1ΔIQ* cells, however we still did not observe any co-localization of Cam2 with Pik1 (Fig. S3A). To examine whether a portion of Cdc4 localized to the trans-Golgi, it was co-imaged with Sec72-mCherry; no colocalization was observed at any cell cycle stage (Fig. S3B). We also combined *pik1-11* with a temperature sensitive allele of *cdc4*, *cdc4-31*, but no genetic interaction was not observed (Fig. S3C). These results are consistent with there being a lack of evidence for a role of Pik1 in cytokinesis. Taken together, our data suggest it is unlikely that Pik1 functions with Cdc4 or Cam2 in either cytokinesis or endocytosis.

Ncs1 is an established Pik1 binding partner and regulator (Lim *et al*., 2011). Therefore, it was not surprising that when we attempted to combine *ncs1Δ* with *pik1-11* we found that they were synthetically lethal (Fig. 4A). Also, as predicted, when Ncs1 was tagged with mCherry, we observed co-localization of Ncs1-mCherry with Pik1-D450-mNG at the trans-Golgi (Fig. 4B). In *S. cerevisiae*, the binding site of the Ncs1 ortholog Frq1 in Pik1 is required for Pik1 Golgi localization implicating Frq1/Ncs1 as necessary for Pik1 Golgi localization (Strahl *et al*., 2005). Because *ncs1* is not an essential gene (Hamasaki-Katagiri et al., 2004), unlike *S. cerevisiae FRQ1* (Hendricks *et al*., 1999), it was possible to test this idea in *S. pombe*. We found that Pik1-D450-mNG still localized to the trans-Golgi marked by Sec72-mCherry in *ncs1Δ* cells (Fig. 4C). Therefore, at least in *S. pombe*, Ncs1 is not required for Pik1 Golgi localization. The small GTPase Arf1 is also implicated in promoting *S. cerevisiae* Pik1 Golgi localization (Highland and Fromme, 2021) so perhaps *S. pombe* relies more on this mechanism. Unfortunately, because *S. pombe* Arf1 is essential and conditional alleles of *arf1* are not available, we were unable to test this possibility. Interestingly, there was a high cytoplasmic Pik1 population in *ncs1Δ* cells that was not observed in wildtype cells; indeed, there was >2-fold more Pik1 overall (Fig. 4D). We currently do not have a mechanistic explanation for this observation.

**Figure 4.**
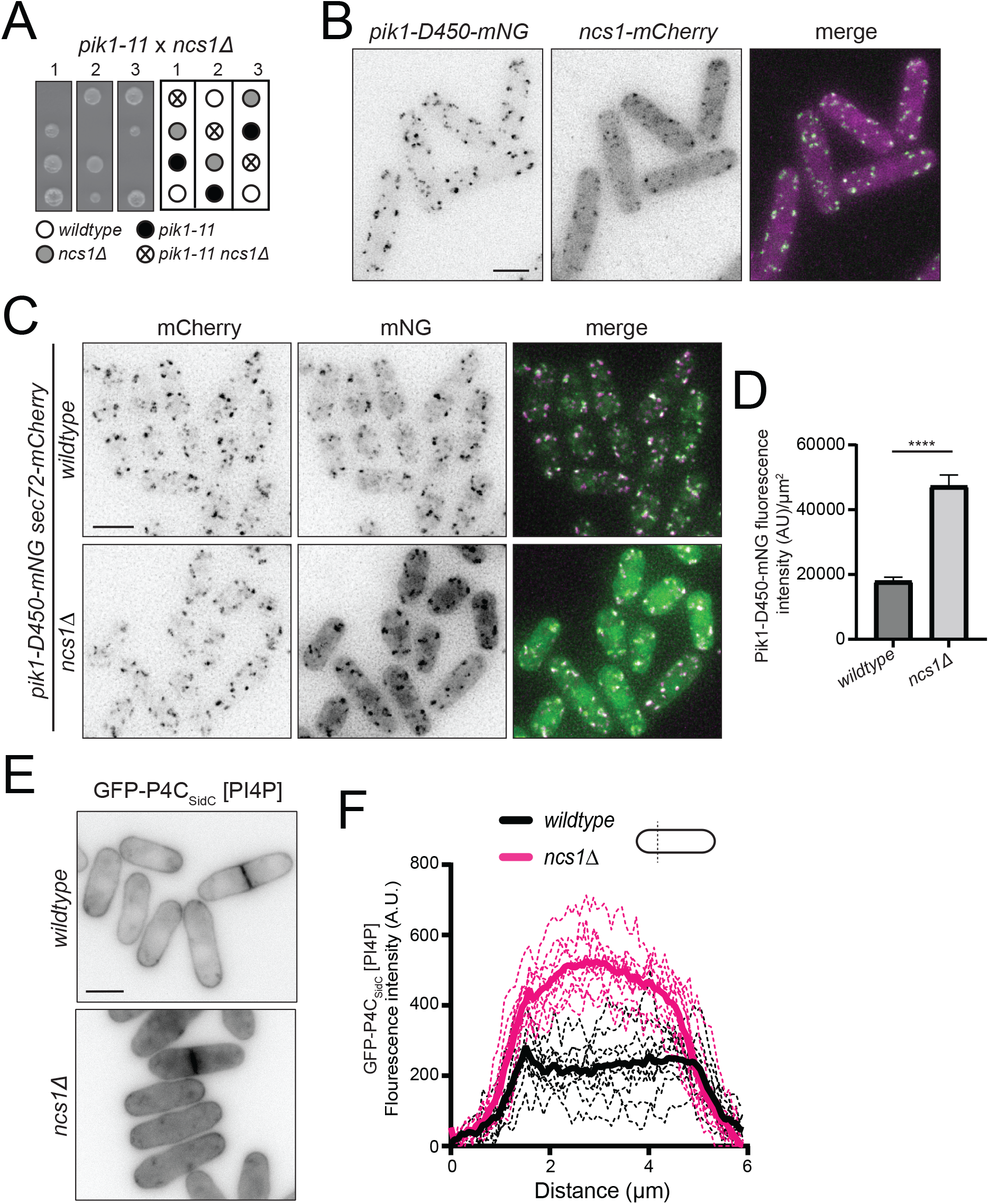
Ncs1 promotes Pik1 function but is not required for Pik1 localization. A) Representative tetrads and schematic of the indicated genetic cross. B) Live-cell imaging of cells expressing Pik1-D450-mNG and Ncs1-mCherry. C) Live-cell imaging of cells expressing Pik1-D450-mNG and Sec72-mCherry in wildtype or *ncs1Δ* cells. D) Quantification of Pik1-D450-mNG whole cell fluorescence intensity from cells in C. n = 15 for each. **** *p* ≤ .0001. Student’s t-test. E) Live-cell images of GFP-P4C_SidC_ in wildtype or *ncs1Δ* cells. F) Line scans of fluorescence intensity drawn across the short axis of 10 cells from E for each strain. Solid lines represent the mean and dotted lines are the individual line scan traces. Scale bars, 5 μm.

The current model for Frq1/Ncs1 function is that it holds the Pik1 kinase domain in an active conformation (Strahl *et al*., 2003, 2005, 2007; Lim *et al*., 2011). To determine whether there was Golgi PI4P in *ncs1*Δ, a proxy for a change in Pik1 activity, we imaged cells expressing GFP-P4C_SidC_ in wildtype and *ncs1Δ* cells. We observed the persistence of Golgi PI4P puncta as well as increased cytoplasmic and PM PI4P levels in *ncs1Δ* cells (Fig. 4E-F). Combined with the fact that Pik1 is essential whereas Ncs1 is not (Hamasaki-Katagiri et al., 2004; Park et al., 2009), it seems unlikely that Ncs1 is required for *S. pombe* Pik1 activity and perhaps even acts as a negative regulator. Future studies will be required to dissect the mechanism of Ncs1-dependent regulation of Pik1 in *S. pombe*.

## Materials and Methods

### Yeast Methods

All *S. pombe* strains used in this study (Table S1) were cultured using standard methods in YE media (Moreno *et al*., 1991; Forsburg and Rhind, 2006). Transformation of yeast with linear DNA was accomplished using a lithium acetate method (Keeney and Boeke, 1994; Forsburg and Rhind, 2006). Strain construction was accomplished through tetrad analysis using standard methods.

Tagged *ncs1* strains were generated by inserting sequences encoding mCherry and natMX6 from a pFA6 cassette the 3′ end of the ORF at the endogenous locus as previously described (Wach *et al*., 1994; Bähler *et al*., 1998). Nourseothiricin (clonNAT, 100 mg/mL, GoldBio; cat# N-500-100) was used for selection of natMX6-containing cells on YE plates. All fusion proteins examined in this study were expressed from their native promoters at their chromosomal loci.

*pik1-D450-mNG* was constructed by cloning synthesized gene blocks (Integrated DNA Technologies) into the PstI site of pIRT2 using Gibson assembly. 300 bp 3’ and 5’ flanks were included as well as the *kanMX6* gene and promoter between the *pik1* stop codon and the 3’ flank. This pIRT2 construct was transformed into cells and G418 (Geneticin, 100 mg/mL, Thermo Fisher Scientific; cat# 11811031) was used to select the appropriate integrants on YE plates. Colonies were confirmed with PCR and imaging.

### Isolation of temperature sensitive alleles with error-prone PCR

The *pik1-11* temperature sensitive allele was constructed as described (Tang *et al*., 2019) with the exception that EX taq polymerase (Takara, cat# 4025) and accompanying dNTPs (Takara, cat# RR01BM) were used.

### Microscopy

Yeast cells were grown at 25°C in YE prior to live-cell imaging unless otherwise grown at 25°C and then shifted to 36°C for 3 hours and imaged at both temperatures. Images were acquired with either 1) a Personal DeltaVision microscope system (Leica Microsystems) that includes an Olympus IX71 microscope, 60X 1.42 NA PlanApo oil immersion objective, a 60X 1.49 TIRF objective, a pco.edge sCMOS camera, and softWoRx imaging software, or 2) with a Zeiss Axio Observer inverted epifluorescence microscope with Zeiss 63X oil (1.46 NA) and captured using Zeiss ZEN 3.0 (Blue edition) software and Axiocam 503 monochrome camera (Zeiss). Images in Figure 2, 4E, S1B and S1D (bottom images) are non-deconvolved maximum intensity projections of 4 medial z sections spaced at 0.5 µm. Images in Figure 2E, 3B, 3D, S1D (top images) and S3 are non-deconvolved maximum intensity projections of z sections spaced at 0.5 µm. Images in Figure 3C and 4 are deconvolved maximum intensity projections of z sections spaced at 0.5 µm. Quantification of images was performed using Fiji (a version of ImageJ software available at https://fiji.sc) (Schindelin *et al*., 2012).

The whole-cell intensity measurement in Figure 4D was corrected for background. In each image used for quantification, background intensity measurements were taken from an area without any cells, which was divided by that area to give the average intensity per pixel of the background. This value was then multiplied by the area of the region of interest (ROI) and subtracted from that ROI’s raw intensity measurement to get the intensity measurement corrected for background. The corrected intensity measurements were divided by the area of the ROI.

For all line scans, a sum projection of 4 medial slices of non-deconvolved images were used. Intensity measurements were plotted against the distance across the short cell axis on cells that were 10-12 μm in length. The line scans were aligned by the first peak in fluorescence intensity.

### Protein structure prediction

Protein structure prediction of Pik1-D450-mNG was generated with the ColabFold interface to the AlphaFold2 pipeline on the Colab platform (AlphaFold2.ipynb) (Jumper *et al*., 2021; Mirdita *et al*., 2022; Varadi *et al*., 2022).

### Statistical analysis

All statistical analyses were performed in Prism 8 (Graphpad software).

## Data availability statement

The data underlying Figures 1-4 and Supplemental Figures 1-3 are openly available in Mendeley Data at doi: 10.17632/tk4bjggsns.1 and 10.17632/t6d5g457rv.1.

## Acknowledgements

We thank Kazutoshi Akizuki and Sierra Cullati for critical reading of the manuscript, Jessie Nguyen for technical assistance and Snezhana Oliferenko for strains. This work was supported by NIH grant R35GM131799 to K.L.G.

## Abbreviations

(mNG): mNeonGreen
(PI): phosphatidylinositol
[PI(4,5)P_2_]: PI-4,5-bisphosphate
(PI4P): PI-4-phosphate
(PIP): phosphoinositide
(PM): plasma membrane
(CR): cytokinetic ring

